# Cross-species communication via *agr* controls phage susceptibility in *Staphylococcus aureus*

**DOI:** 10.1101/2023.04.03.535347

**Authors:** Jingxian Yang, Janine Zara Bowring, Janes Krusche, Benjamin Svejdal Bejder, Stephanie Fulaz Silva, Martin Saxtorph Bojer, Tom Grunert, Andreas Peschel, Hanne Ingmer

## Abstract

Bacteria and their viruses (phages) use quorum sensing (QS) systems to coordinate group behavior. In *Staphylococcus aureus*, QS plays a critical role in the transition from colonization to infection and involves the accumulation of auto-inducing peptides (AIPs). Humans and animals are also colonized by non-aureus staphylococci (NAS) that produce AIPs, many of which inhibit *S. aureus* QS. We found that QS induction is necessary for *S. aureus* susceptibility to the lytic phage, Stab20 and that in mixed communities with NAS producing inhibitory AIPs, *S. aureus* is protected from phage infection. The primary phage receptors in *S. aureus* are wall teichoic acids (WTA) substituted with α- and/or β-linked N- acetylglucosamine (GlcNAc). We show that QS induction reduces α-GlcNAc substitutions and enables Stab20 infection through binding to β-glycosylated WTA. However, in the presence of inhibitory AIPs or during co-culture with NAS, QS induction and Stab20 infection are impeded. Our results highlight how cross-species communication can significantly impact bacterial susceptibility to phages and may explain occasional failures observed when phages are used as antimicrobials in for example phage therapy.

## Introduction

Bacterial viruses, the bacteriophages (or phages), have received renewed interest as antimicrobial options for treating infections with antibiotic resistant pathogens. Phages are classified as lytic or temperate with the former infecting and killing bacteria while the latter integrate into and replicate with the bacterial genome (Abedon, 2005). In phage therapy, lytic phages are employed to eradicate infections and historically, there are examples of life- threatening infections being cured with phages (Hatfull *et al*, 2022). Yet, phage therapy can fail for example due to phage resistance (Pires *et al*, 2020).

One of the pathogens for which phage therapy has been applied is *Staphylococcus aureus* (Petrovic Fabijan *et al*, 2020; Ramirez-Sanchez *et al*, 2021). It is an opportunistic pathogen that on the one hand colonizes a third of the human population, asymptomatically, but also can cause a variety of serious infections. Transition from colonization to infection is in part controlled by the regulatory *agr* quorum sensing (QS) system composed of a two-component sensory histidine kinase and response regulator as well as the auto-inducing, cyclic peptides (AIPs). In response to cell density and AIP concentration, *agr* mediates the transition from expression of *S. aureus* adhesion and colonization factors at low cell densities to expression of a large array of toxins and extracellular enzymes at high cell densities (Novick & Geisinger, 2008). Curiously, *S. aureus* strains encode one of four *agr* variants (I-IV), where a cognate AIP is required for *agr* induction while some non-cognate AIPs prevent *agr* induction. AIPs are also produced by a variety of non-aureus staphylococci (NAS) that colonize humans and animals and a great part of these interfere with induction of *S. aureus* QS, such as those produced by *S. haemolyticus, S. caprae* and *S. pseudintermedius* (Canovas *et al*, 2016; Gless *et al*, 2021; Horswill & Gordon, 2020; Peng *et al*, 2019b). These interactions are likely to be key for both *S. aureus* colonization and infection (Baldry *et al*, 2018; Nakamura *et al*, 2020; Paharik *et al*, 2017; Parlet *et al*, 2019).

The main receptors for *S. aureus* phages are the wall teichoic acid (WTA) glycopolymers that decorate the cell surface by being attached to the cell wall (Xia *et al*, 2011). WTA can be modified by alanylation and glycosylation with the latter being important for interactions with the innate and adaptive immune systems as well as for phage susceptibility (van Dalen *et al*, 2020; Weidenmaier & Peschel, 2008). For example, most myophages recognize the WTA backbone regardless of glycosylation (Peng *et al*, 2019a; Xia *et al*., 2011) whereas, α- or β-glycosylated WTA is essential for the infection by siphophages and podophages (Li *et al*, 2015; Li *et al*, 2016; Xia *et al*., 2011; Xia *et al*, 2010), as well as for a few myophages (Azam *et al*, 2018; Estrella *et al*, 2016).

WTA synthesis involves initiation (*tarO*), priming (*tarABDF*) and polymerization (*tarIJLK*) of repeating ribitol phosphate (RboP) units and glycosylation is catalyzed by the TarM and TarS that add N-acetylglucosamine (GlcNAc) to the C4 position of the RboP unit in an α- or β-configuration, respectively (Winstel *et al*, 2015; Xia *et al*., 2010). In addition, some phages encode TarP, which modifies the C3 position with a β-O-linkage (Gerlach *et al*, 2018). After synthesis, the WTA is translocated to the cell surface by the ABC transporter, TarGH, where it is further decorated with D-alanine and anchored to the peptidoglycan (Meredith *et al*, 2008; Winstel *et al*., 2015).

Here we have examined how *S. aureus agr* affects phage susceptibility and we find that successful infection by the lytic myophage, Stab20 relies on *agr* induction, resulting from decreased expression of α-glycosylation of the WTA. To our knowledge, this is the first report to show that *agr* controls phage infections and alters WTA glycosylation patterns. Further we show that in mixed communities with NAS producing inhibitory AIPs, *S. aureus* is protected from Stab20 infection. Thus, our findings indicate that there is a delicate balance between the ubiquitous NAS and *S. aureus* that impacts not only *S. aureus* virulence factor production but also WTA-mediated immune responses, phage susceptibility and with that the composition of entire microbial communities. In perspective, our results suggest that the composition of bacterial communities may greatly influence how susceptible bacteria are to phages and thus, may explain why phage therapy occasionally fails.

## Results

### Induction of *agr* promotes phage infection of *S. aureus*

To explore if the *S. aureus agr* QS system affects susceptibility to phage infections, we examined a collection of phages including temperate phages belonging to serogroup B (ɸ11, ɸNM1, ɸ80 and ɸ52A), serogroup Fb (ɸ13) and serogroup A (ɸ12, ɸ42b and ɸ42f) and lytic phages (ɸIPLA-RODI, Stab20 and Stab21) belonging to serogroup D. The phages were used to infect an *agrA* deletion mutant strain constructed in TB4 (wild type), a prophage-cured *S. aureus* derivative of strain Newman (Table S1), as well as TB4 cells treated with synthesized and purified AIP-I inducing *agr* or the AIP from *S. hyicus* (AIP_hy_) that inhibits *S. aureus* QS (Gless *et al*., 2021). In comparison to untreated WT cells, we found that addition of AIP-I increased susceptibility to a variety of both lytic and temperate phages, namely ɸIPLA-RODI, Stab20, Stab21, ɸ13, ɸ12, ɸ42b, ɸ42f and ɸ80, with the greatest effect being on Stab20 infection. For Stab20, we only observed infection when cells were grown in the presence of AIP-I prior to phage infection, whereas no infection was observed for untreated cells, cells lacking *agrA*, or cells treated with the inhibitory AIP_hy_ (Fig S1). The effect of AIP-I on *S. aureus* susceptibility to Stab20 infection was further confirmed in a liquid infection assay where only AIP-I treated WT cells were killed by the phage in contrast to untreated WT cells, *agrA* mutant cells or cells treated with AIP_hy_ (Fig S2). Thus, we decided to focus our studies on Stab20.

**Table 1.** Stab20 infection at different growth phases determined as plaque forming units (PFU).

| Strains | Phage Titer (PFU/mL) <sup>a</sup> |  |  |
| --- | --- | --- | --- |
|  | Exponential (OD<sub>600</sub> 0.35) | Early stationary (OD<sub>600</sub> 1.4) | Late stationary (overnight) |
| TB4+AIP-I | ( 0.9 ± 0.07 ) *10<sup>8</sup> | ( 1.3 ± 0.07 ) *10<sup>8</sup> | ( 1.4 ± 0.09 ) *10<sup>8</sup> |
| TB4 | -<sup>b</sup> | - | ( 1.4 ± 0.05 ) *10<sup>8</sup> |
| TB4+AIP<sub>hy</sub> | - | - | - |
| TB4Δ*agrA*+AIP-I | - | - | - |
| TB4Δ*agrA* | - | - | - |

a: phage titer was calculated by the full plate plaque assay.
b: no visible plaques.

Stab20 is a lytic myophage within the genus Kayvirus in the subfamily Twortvirinae (Oduor *et al*, 2019). It was isolated on *S. xylosus* and has a broad host range infecting both methicillin resistant (MRSA) and susceptible *S. aureus* (MSSA) as well as some strains of *S. epidermidis, S*. *haemolyticus* and *S. saprophyticus* (Oduor *et al*, 2020). Given that infection of *S. aureus* by Stab20 was stimulated by AIP-I, we examined how susceptibility varied with growth phase. As expected, AIP-I promoted Stab20 infection of WT cells in both exponential and stationary growth phase, whereas in the absence of exogenous AIP-I it only infected cells in the stationary growth phase (Table 1). In contrast, phage infection was abolished in stationary phase cells treated with the inhibitory AIP_hy,_ and in the Δ*agrA* mutant cells treated with AIP-I. These results corroborate that induction of *agr* controls susceptibility to Stab20 infection.

To determine whether *agr* impacts Stab20 infection in other *S. aureus* strains than TB4 we examined strains belonging to different clonal complexes and *agr* types, namely CC8 (TB4, JE2, NCTC8325, COL and 110900), CC398 (55-100-002), CC5 (SA564, Mu50 and N315), CC1 (MW2) and CC30 (UAMS-1 and MN8). The cells were grown either in the presence or absence of their cognate AIPs (Table S2) and were subsequently infected either at exponential or stationary growth phase with Stab20 (Fig 1). The results show that several strains were infected independently of growth phase and AIP-addition, including COL, 110900, SA564 and MN8, while some were not infected at all (NCTC8325, 55-100-002 and N315). MW2 resembled TB4 in being infected upon AIP induction and in stationary growth phase, while JE2 and UAMS-1 were infected only when being in stationary phase and exposed to inducing AIPs. Thus, infection by Stab20 is strain specific and is not associated with the clonal complex or the AIP type of the strains.

**Figure 1.**
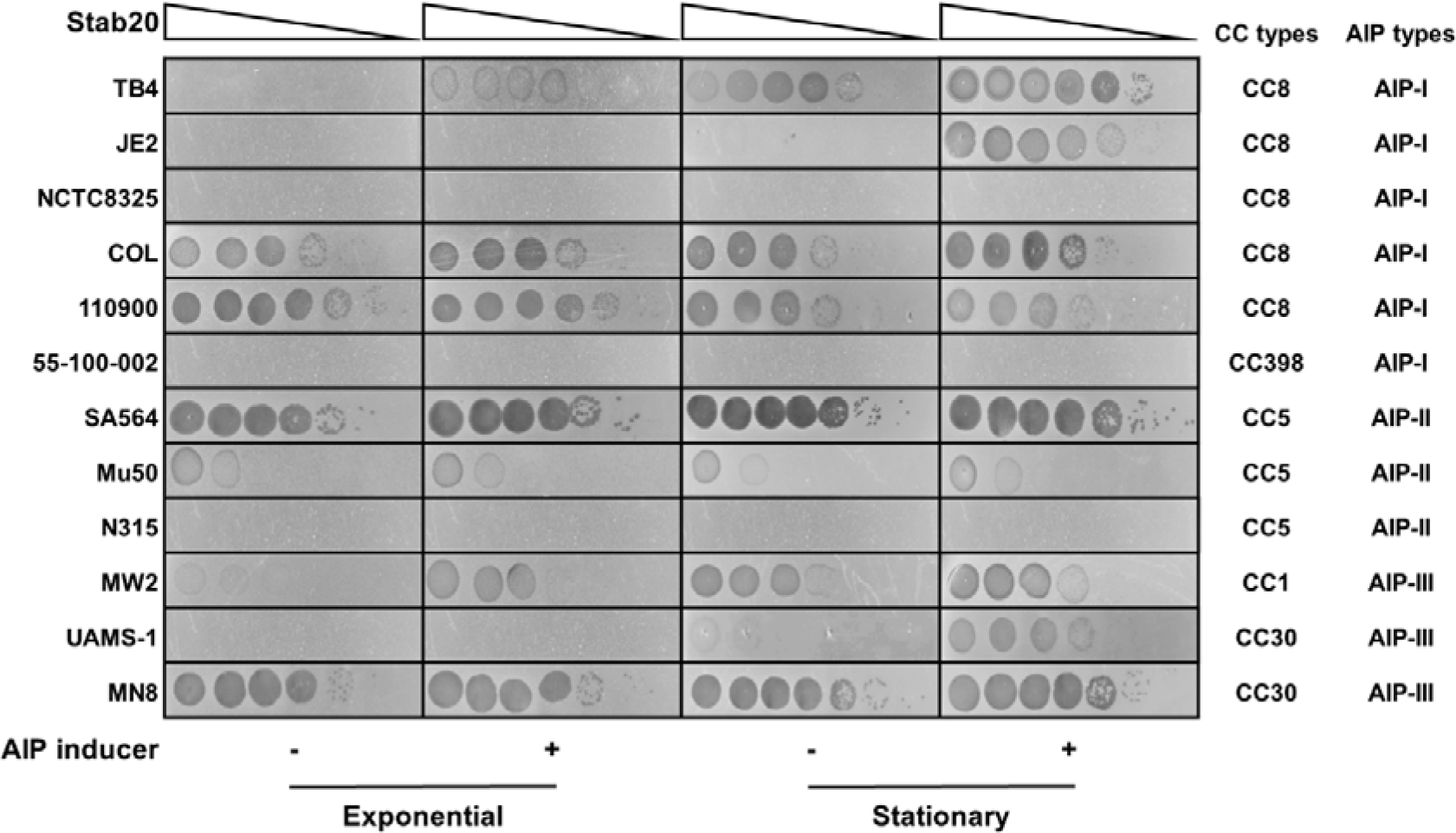
Infection of *S. aureus* strains by Stab20. Stab20 phage lysate dilutions (10^0^∼10^-6^) were spotted onto plates carrying cells grown either with (0.1 µM) or without AIP inducer and harvested in either exponential or stationary growth phase. JE2, NCTC8325, COL, 55- 100-002, and 110900 were induced by AIP-I, SA564, Mu50 and N315 were induced by AIP- II, MW2, UAMS-1 and MN8 were induced by AIP-III.

### Glycosylation of wall teichoic acids affects Stab20 infection

WTA is the primary receptor of staphylococcal phages and therefore, we speculated that *agr* may affect WTA synthesis or structure. To test this, we infected strain JE2 and its Δ*tarO*, *tarM*, *tarS* and *tarK* mutant derivatives with Stab20 and examined plaque formation (Fig 2). Here we observed that deletion of *tarO,* which abolishes WTA entirely, prevented Stab20 infection as has been observed for other phages (Li *et al*., 2015; Xia *et al*., 2011). Inactivation of *tarK* did not affect phage susceptibility while inactivation of *tarS* abolished phage infection in stationary phase cells supplemented with AIP-I. The greatest effect, however, was seen for *tarM* mutant cells that were infected by Stab20 independently of growth phase and AIP addition. These results suggest that *tarS* is required for phage infection and that both *tarM* and *tarS* expression maybe affected by *agr*.

**Figure 2.**
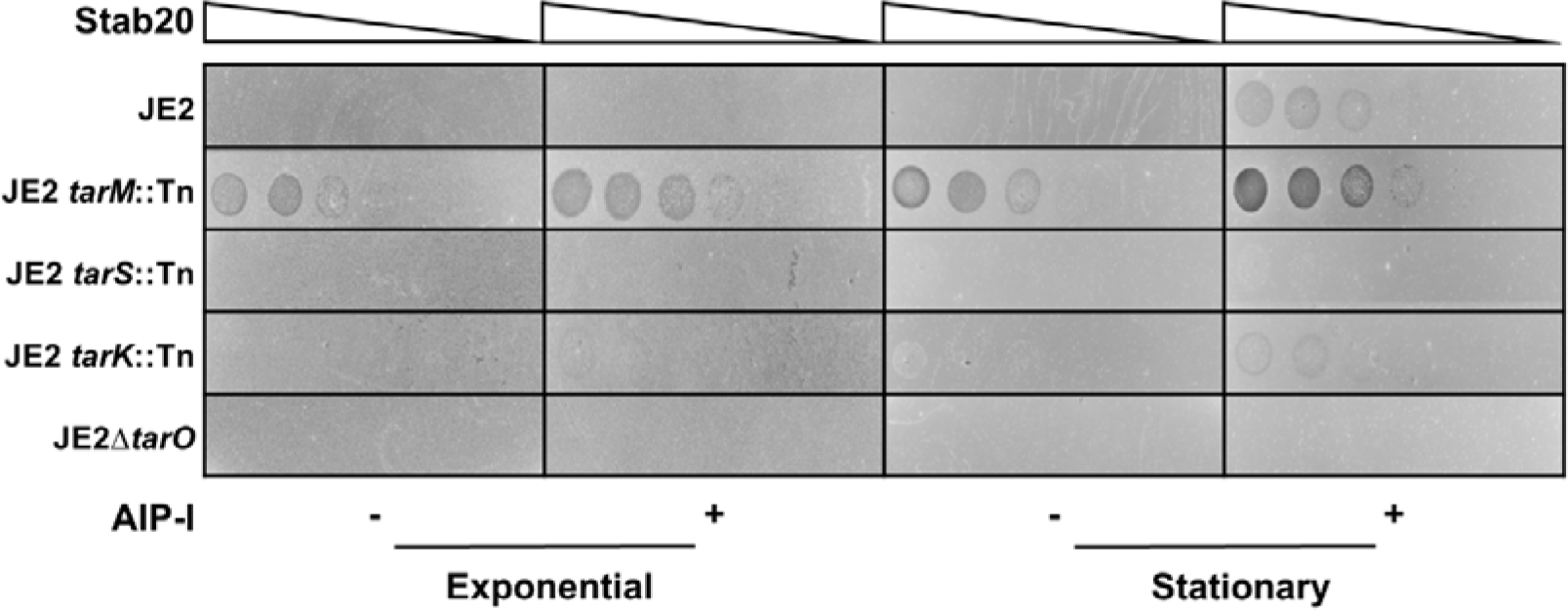
Stab20 infection of WTA mutant derivatives. Dilutions of Stab20 phage lysate (10^0^∼10^-6^) were spotted onto plates carrying JE2 or mutant derivatives defective in WTA grown either with (0.1 µM) or without AIP-I and harvested in either exponential or stationary growth phase.

### *agr* downregulates *tarM* expression

As our data suggested that *tarM* and/or *tarS* could be *agr* regulated, we monitored gene expression in TB4 WT cells either with or without inducing AIP-I as well as in *agrA* mutant cells. Here we observed that transcription of *tarM* decreased significantly (*p*<0.05) in cells treated with the inducing AIP-I while there were no significant differences in *tarS* expression (Fig 3). This result agrees with a previous study identifying genes in strain MW2 being down-regulated by *agr* with MW0919 corresponding to *tarM* (Queck *et al*., 2008).

**Figure 3.**
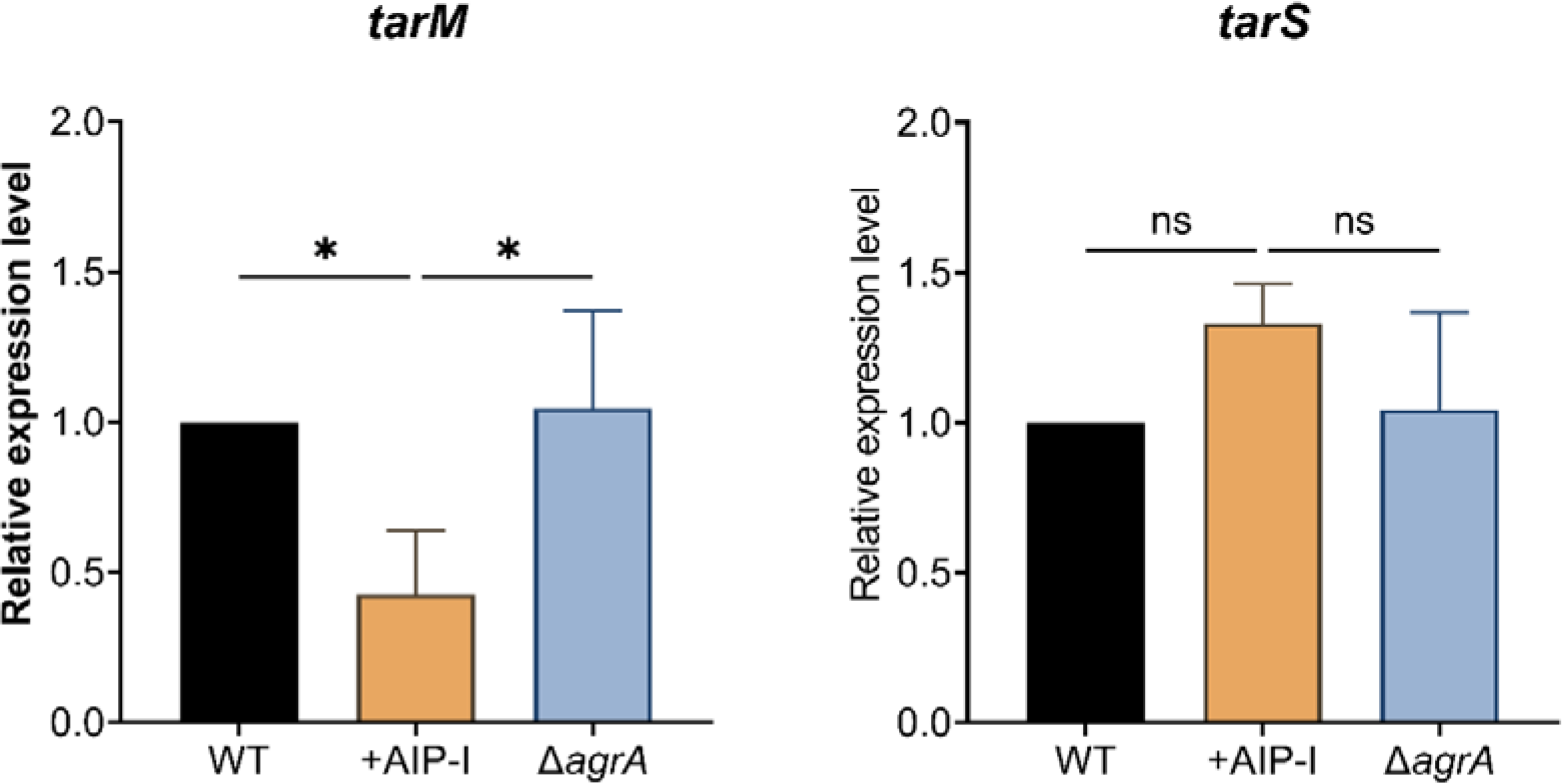
Expression of *tarM* is *agr* regulated. Relative expression levels of *tarS* and *tarM* in TB4, AIP-I-treated TB4 (0.1µM), and TB4Δ*agrA* at OD_600_ of 0.35 were determined by RT- qPCR using *pta* as a reference gene. Data are shown as the means with standard deviation from three biological replicates. Data were analyzed using one-way ANOVA. \**P* < 0.05; \*\**P* < 0.01; and \*\*\**P* < 0.001.

To confirm our results, we deleted *tarS* and *tarM* in TB4 WT and Δ*agrA* mutant cells and examined Stab20 susceptibility. Similar to the observation in JE2, deletion of *tarM* in TB4 allowed Stab20 to infect independently of growth phase and AIP-I addition, and even allowed infection of *agrA* mutant cells. Importantly, the phenotype of the *tarM* mutant was complemented by ectopic expression of *tarM*. In contrast, Stab20 was unable to infect *tarS* mutant cells in stationary phase while infection was restored by a plasmid-encoded *tarS*. When deleting both *tarM* and *tarS,* Stab20 infection was prevented entirely (Fig 4). These results are consistent with a model where the β-GlcNAc decoration of WTA catalyzed by

**Figure 4.**
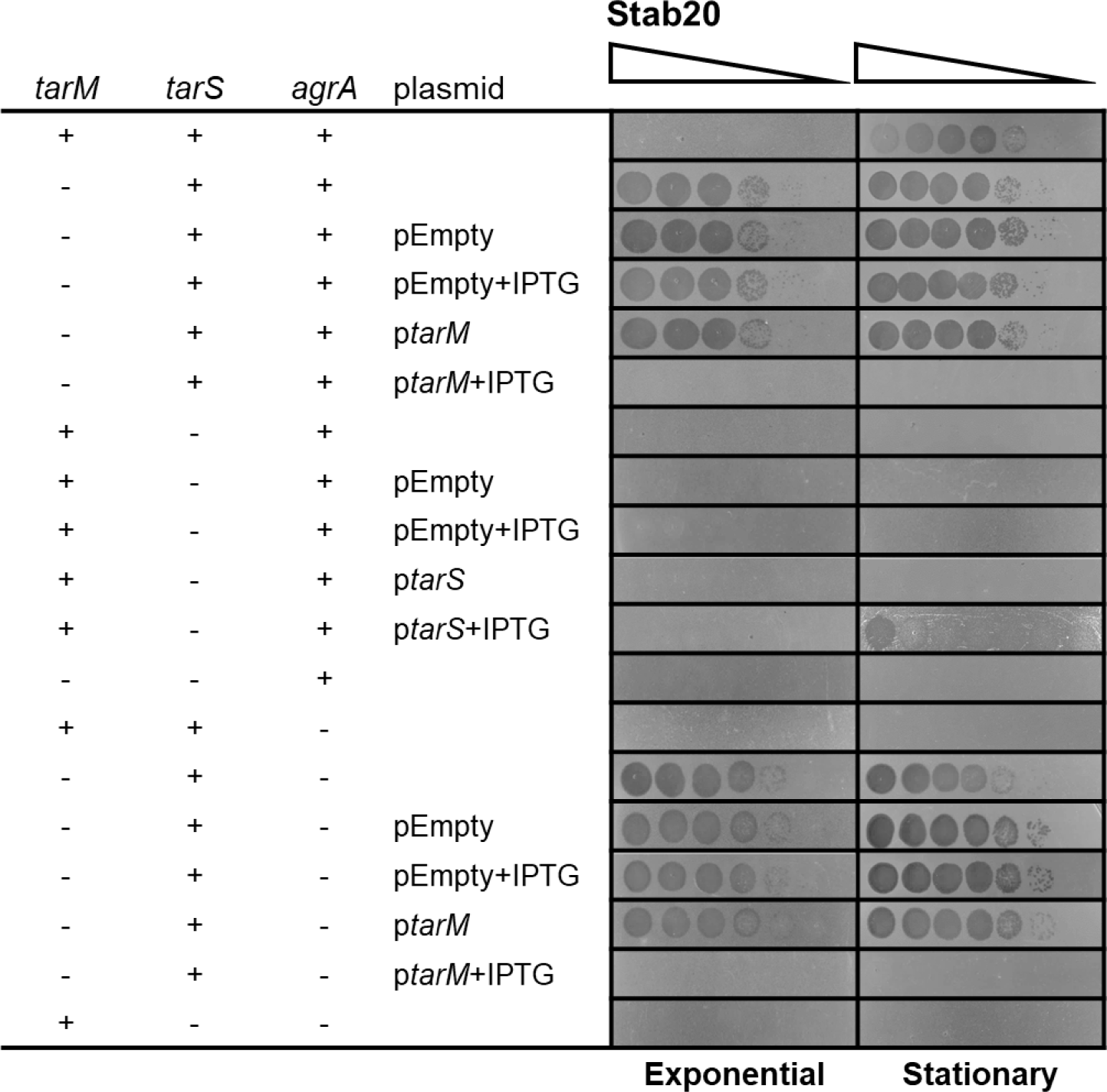
Stab20 infects *tarM* but not *tarS* mutant cells. Deletion mutants of *tarM*, *tarS* and both *tarM* and *tarS* in TB4 or TB4Δ*agrA* as well as their complemented derivatives were infected with Stab20 (10^0^∼10^-6^). IPTG was added at 400µM.

TarS is the receptor of Stab20 while the α-GlcNAc decoration catalyzed by TarM interferes with phage infection and is repressed upon *agr* induction.

To further monitor the impact of *agr* on Stab20 infectivity, we expressed *tarM* from an IPTG-inducible promoter in pSK9067 and examined infection at various IPTG concentrations (Fig 5). In the absence of IPTG, Stab20 eliminated growth of AIP-I treated cells but did not affect untreated cells. However, with increasing concentrations of IPTG and thus increased expression of *tarM,* phage killing was delayed and even prevented at 400 μM IPTG. These results show that TarM is key in preventing Stab20 infection and acts in a concentration-dependent manner.

**Figure 5.**
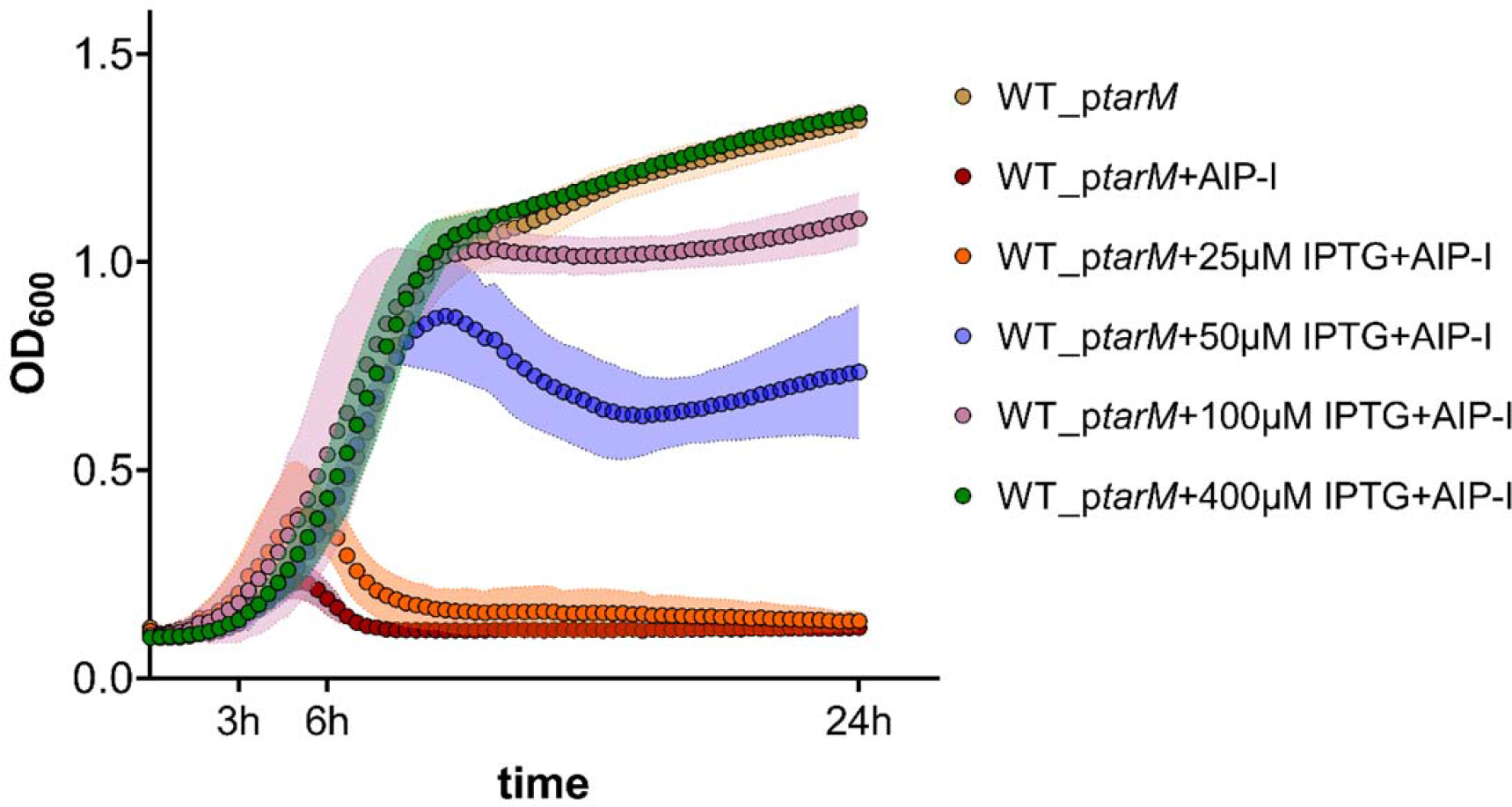
Induction of *tarM* expression abolished Stab20 infection. Growth of TB4_p*tarM* treated with (0.1µM) or without AIP-I upon Stab20 infection. Expression of *tarM* was induced by IPTG at indicated concentrations and infection occurred at a multiplicity of infection (MOI) of 0.1. Data are represented as mean ± SD with three biological replicates.

In light of the observation that *agr* promotes phage infection by suppressing *tarM* expression, we examined the level of α-GlcNAc modification of WTA by reacting WT and AIP-I treated cells as well as the Δ*tarM* and Δ*tarS* mutants with anti-α-GlcNAc-WTA IgG and monitoring binding by flow cytometry (Hendriks *et al*, 2021). As predicted, this analysis showed that the Δ*tarM* and the Δ*tarM*Δ*tarS* double mutants had no α-GlcNAc substitutions on their WTA, whereas the Δ*tarS* mutant still had α-GlcNAc substitutions comparable to the TB4 WT cells. Remarkably, the amount of α-GlcNAc was dramatically decreased when cells were treated with inducing AIP-I, showing that activation of *agr* reduces α-GlcNAc modification of WTA (Fig 6).

**Figure 6.**
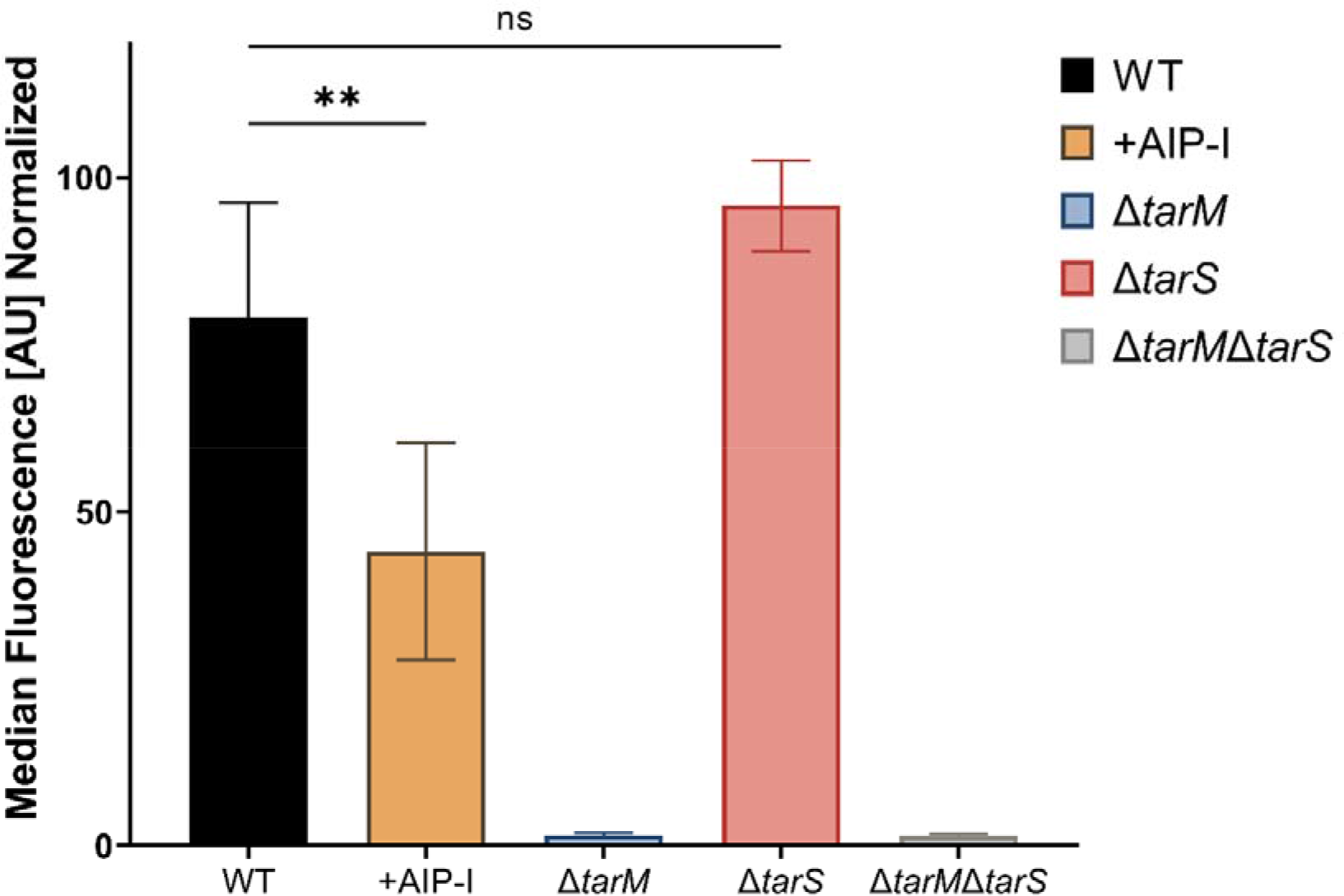
*agr* induction reduced α-GlcNAc substitutions on WTA. WT, WT treated with inducing AIP-I (1µM), Δ*tarM*, Δ*tarS* and Δ*tarM*Δ*tarS* were tested for binding to IgG targeting α-GlcNAc-WTA. Data are shown as the means with standard deviation from three biological replicates. Data were analyzed using one-way ANOVA. \**P* < 0.05; \*\**P* < 0.01; and \*\*\**P* < 0.001.

### Non-aureus staphylococci inhibit *agr* and protect against Stab20

As AIPs from non-aureus staphylococci (NAS) strongly influenced the ability of Stab20 to infect *S. aureus*, we examined how co-cultures of NAS together with *S. aureus* influenced Stab20 infectivity of *S*. *aureus* within the combined populations (Fig 7A). We examined *S. caprae, S. haemolyticus* and two strains of *S. pseudintermedius* that all produce AIPs inhibiting *S. aureus agr* (Canovas *et al*., 2016; Gless *et al*, 2019) and that were not infected individually by Stab20 (Fig 7B). Importantly, when co-cultured with TB4 they all prevented or strongly inhibited Stab20 infection (Fig 7B) and the effect was not due to growth inhibition, as TB4 formed the greater part of the co-cultures (Table S4). However, in the absence of *tarM*, the NAS strains were unable to prevent Stab20 infection and also deletion of the *agr* locus in *S. pseudintermedius* ED99 eliminated the inhibitory effect of the strain on phage infection (Fig 7B). Thus, our results show that *agr* mediated repression of *tarM* expression allows phage infection in stationary growth phase and that this effect is mitigated in mixed communities with NAS producing inhibitory AIPs.

**Figure 7.**
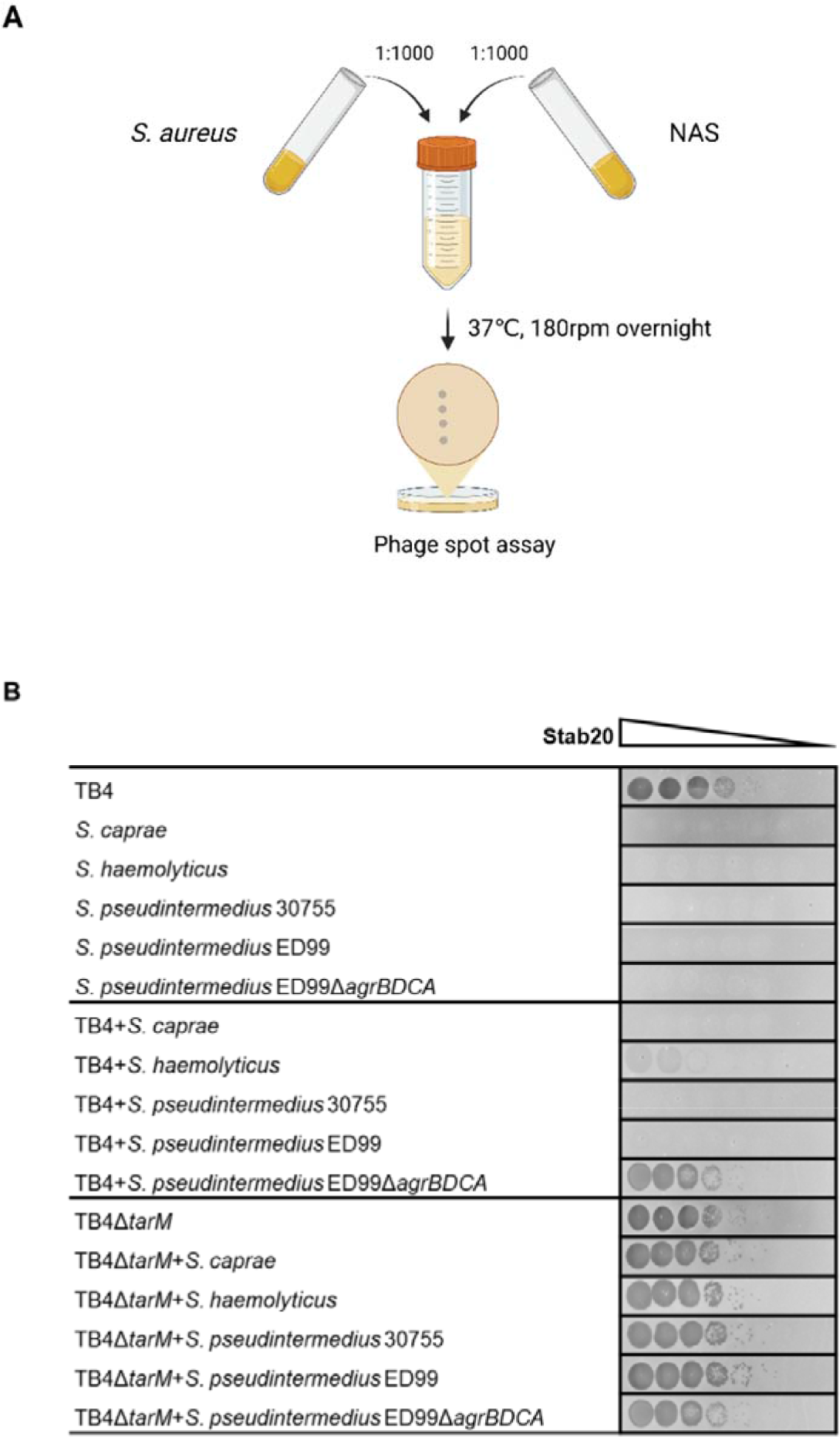
Non-aureus staphylococci inhibit Stab20 infection of *S. aureus*. A. The co-culture was set by diluting standardized overnight culture of tested strains to TSB (1:1000) and overnight co-cultures were collected to measure phage susceptibilities by spotting phage lysate dilutions on the bacterial lawn; B. Susceptibilities of monocultures or co-cultures of TB4 or TB4Δ*tarM* with non-aureus *Staphylococcus* strains.

## Discussion

A key finding of this study is that the non-aureus staphylococci (NAS) prevent phage infection of *S. aureus* by producing variant AIPs that inhibit the *S. aureus agr* and reduce α- GlcNAc glycosylation of WTA, as summarized in Fig. 8. Staphylococci are part of the common skin flora of humans and animals with *S. aureus* being the most prominent opportunistic pathogen. On the skin surface, *S. aureus* exists in tightly interwoven microbial communities (Grice & Segre, 2011) where it co-colonizes with NAS such as *S. haemolyticus* and *S. caprae* (Cosseau *et al*, 2016) and for people in contact with animals also *S. pseudintermedius* (Somayaji *et al*, 2016). In addition to NAS, a wide range of other bacterial species produce molecules that prevent *S. aureus agr* activation. For example, Lactobacilli and *Bacillus subtilis* produce cyclic peptides different from AIPs that in the case of fengycin is active in the human gut (Li *et al*, 2011; Piewngam *et al*, 2018) while multiple commensal *Corynebacterium* spp. produce yet unidentified compounds that also inhibit *S. aureus agr* (Ramsey *et al*, 2016). When these different bacteria share an ecological niche, intricate interactions are possible via AIP-mediated signaling cross-talk that may impact both *S. aureus* colonization and infection (Baldry *et al*., 2018; Nakamura *et al*., 2020; Paharik *et al*., 2017; Parlet *et al*., 2019; Peng *et al*., 2019b; Williams *et al*, 2019). Our results show that they can also impact whether *S. aureus* is infected or not by lytic phages, which in turn will influence microbial community composition and susceptibility to phage therapy.

**Figure 8.**
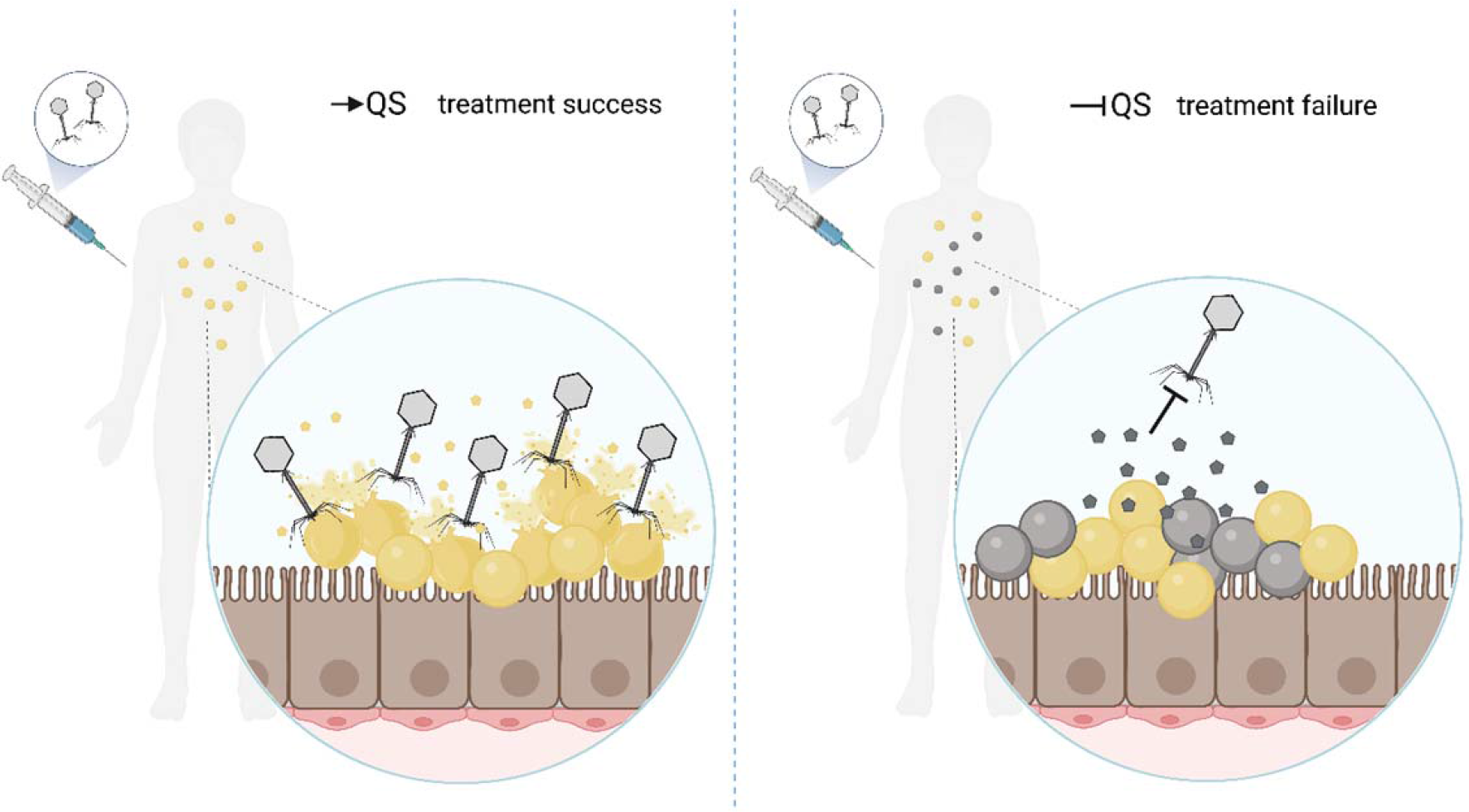
Impact of *agr*-mediated interspecies communication on phage therapy. Hypothetical scenarios of staphylococcal infection caused by mono-*S. aureus* (left) or mixed staphylococci (right), respectively. *S. aureus* (yellow) and NAS (grey) are represented by the spheres in each panel and the associated AIPs are shown by the pentagons.

Recently, reports have documented that phages themselves also respond to environmental signals via QS. In *Vibrio cholera* and *Bacillus subtilis,* temperate phages employ QS to make decisions on lysis-lysogeny either by tapping into the host QS systems or by encoding an “arbitrium” sensing system (Erez *et al*, 2017; Silpe & Bassler, 2019a, b), while in *Pseudomonas aeruginosa,* the bacterial QS systems stimulate susceptibility to at least two lytic phages (Broniewski *et al*, 2021). Bacterial cross-talk was also shown to impact the biology of a temperate *V. cholera* phage where a phage encoded regulator determines whether the phage lifestyle is lytic or lysogenic (Silpe *et al*, 2022). Thus, multi-species bacterial interactions appear to impact phage biology in both Gram-positive and -negative bacteria.

In staphylococci, the structure of WTA is diverse and the glycoepitope(s) can vary between strains and clones (Du *et al*, 2021). In *S. aureus*, three forms of WTA glycosylation have been reported (Tamminga *et al*, 2022) and variation in the corresponding glycosyltransferases determines the host range of staphylococcal phages (Moller *et al*, 2019). The twelve *S. aureus* strains examined here all had *tarS*, 7 harbored *tarM* and 3 encoded *tarP* (Table S2). Although *tarS*-mediated β-GlcNAc modification was found to be the receptor of Stab20 (Fig 4), the *S. aureus* strains differed in their susceptibility as strains NCTC8325, 55-100-002 and N315 had *tarS* but failed to be infected by the phage (Fig 2, Table S2). Intriguingly, the presence of *tarM* appears to be inversely associated with phage susceptibility, as five of seven *tarM*-negative strains were susceptible to Stab20, and the two unsusceptible strains also carried *tarP* (55-100-002 and N315) (Table S2). This is consistent with earlier studies on podophages and myophages; *S. aureus* strains lacking *tarM* were all susceptible to podophages Φ44AHJD, Φ66, ΦP68, except those also lacking *tarS* (Estrella *et al*., 2016; Li *et al*., 2015). Notably, prophage-encoded *tarP* might impair phage infection in *S. aureus* since *tarP*-carrying 55-100-002 and N315 showed phage resistance even though *tarM* was absent on their genomes (Table S2).

Glycosylation of the WTA can also be affected by environmental conditions. For example, for *S. aureus* strain Newman it was reported that under *in vitro* conditions it produces almost exclusively α-GlcNAc WTA (>90%), whereas an *in vivo* in an infection the WTA structure is switched to β-GlcNAc (Mistretta *et al*, 2019). Our finding that induction of *agr* downregulates *α*-GlcNAc modifications on WTA in TB4, a strain derived from Newman, strongly suggests that *agr* may be responsible for such a shift in glycosylation patterns. In addition to being a phage receptor, the WTA is important for *S. aureus* in a number of other aspects including pathogenicity, antibiotic resistance, adhesion, colonization and immune interactions with host cells (Brown *et al*, 2012; Guo *et al*, 2021; Ingmer *et al*, 2019; van Dalen *et al*., 2020; Wanner *et al*, 2017; Xia *et al*., 2011; Zhu *et al*, 2018). Thus, the *agr* mediated control of *tarM* may have wide impact on staphylococcal biology beyond interactions with phages.

Our finding that bacterial susceptibility to phages depends on the composition of the community in which they reside has several perspectives. When developing phage therapy, phages are generally characterized by infections of individual strains but not of bacterial communities. Therefore, failures in therapy may arise from bacteria being present in an environmental setting where phage receptors are being masked or their expression repressed and thus, they are resilient to infection (Fig. 8). Another perspective is that environmental regulation of phage receptors may allow for coexistence of lytic phages and theirs hosts, where a fraction of the host population express receptors while the remaining do not. Such phenotypic heterogeneity in phage susceptibility has been seen for *Salmonella enterica* with phase variation in expression of the O-antigen phage receptor (Cota *et al*, 2015). Epigenetic regulation of phage receptors is likely to be much more prevalent than currently recognized and future studies should be directed at understanding how phages interact with their hosts when present in multi-species and natural environments.

## Materials and Methods

### Bacteria and phages

Bacterial strains used in this study are described in Table S1. *S. aureus* strains were cultured in tryptic soy broth (TSB) or tryptic soy agar (TSA), and *E. coli* strains were grown in Luria– Bertani (LB) or LB agar (LA). Antibiotics (erythromycin 5µg/mL or 10µg/mL; ampicillin 100µg/mL; chloramphenicol 10µg/mL) were added as required. Isopropyl β-D-1- thiogalactopyranoside (IPTG; Thermo Scientific) was used for induction of gene expression and X-gal (X-Gal 5-Bromo-4-chloro-3-indolyl-b-D-galactopyranoside; Thermofisher) TSA plates were used for bacterial assessment. AIP-I/II/III and AIP_hy_ were synthesized as described (Gless *et al*, 2017). To induce or inhibit the *agr* system in *S. aureus* strains, overnight cultures were diluted to OD_600_ of 0.01 and grown with the addition of 0.1 μM AIP inducers or AIP_hy_.

### Phage induction and infection

Lytic phage propagation was performed as described (Bowring *et al*, 2022). Briefly, overnight cultures of recipient strains were grown to OD_600_ of 0.15, collected by centrifugation, and resuspended in 1:1 TSB and phage buffer (PHB; MgSO_4_ 1 mM, CaCl_2_ 4 mM, Tris-HCl pH 8.0 50 mM, NaCl_2_ 0.1 M). Suspended cells were infected with a phage stock at an appropriate multiplicity of infection before incubated at 30°C, 80 rpm to a complete lysis. Lysates were filtered and stored at 4°C. Phage induction was performed as described (Bowring *et al*, 2022). Briefly, lysogens were grown overnight and were subcultured to OD_600_ of 0.15, followed by addition of 2 μg/mL mitomycin C (Sigma). Cell cultures were then incubated at 30°C, 80 rpm until completely lysed. Lysates were filtered and stored at 4°C.

### Phage susceptibility assay

Phage full plate plaque assay was conducted by diluting overnight cultures of *S. aureus* to OD_600_ of 0.01 with TSB and grown to either exponential phase (OD_600_ of 0.35) or overnight. Phage lysates were serial diluted in PHB. 100μL recipient cells and 100 μL phage lysates were mixed and incubated for 10 min at room temperature, followed by the addition of 3 mL phage top agar (PTA; 4 g Nutrient Broth No. 2; 0.7g agar), and plated out on phage base agar plates (PBA; 18 g of Nutrient Broth No. 2; 6.3g agar) supplemented with CaCl_2_ (final concentration 10mM). The number of phage plaques was counted after 24h incubation at 37°C. Phage spot assay was performed by spotting 10μL phage lysates onto PBA plates overlaying with 3mL PTA carrying 100 μL recipient cells. Images of phage plaques were obtained by scanning the plates after 24h incubation at 37°C.

Phage liquid infection was assessed in a plate reader (Bioscreen C, Oy Growth Curves Ab Ltd) where cultures grown until OD_600_ of 0.15 (with AIPs or IPTG if necessary) were transferred to honeycomb Bioscreen plates (95025BIO) in 125 μL aliquots. The same volume of phage lysates in PHB were added to each well and OD_600_ was measured every 20 mins for 24 hours at 30°C with shaking.

### RT-qPCR

For RNA sample preparation, overnight cultures were diluted to OD_600_ 0.01 in fresh medium and supplemented by 0.1 µM AIP-I (if necessary). Cells were collected at OD_600_0.35 for RNA isolation by using the RNeasy kit (Qiagen). PrimeScript™ RT Reagent Kit (TAKARA) was used to generate cDNA and FastStart Essential DNA Green Master (Roche) for qPCR in a Lightcycler 96 (Roche). All RT-qPCR experiments were performed in triplicate with five technical replicates, primers are listed in Table S3. Gene *pta* was set as control. Data analysis was performed in the LightCycler Application Software, version 1.1 (Roche).

### Gene cloning and mutant construction

Plasmids (Table S1) and primers (Table S3) used for gene cloning are listed. TB4Δ*agrA* was constructed as previously described (Chen *et al*, 2017). TB4Δ*tarM*, TB4Δ*agrA*Δ*tarM*, and TB4Δ*tarM*Δ*tarS* mutants were constructed by using the SLiCE cloning method (Monk & Stinear, 2021). Primer pairs MA/MB (or SA/SB) and MC/MD (or SA/SD) were used for amplifying upstream and downstream of *tarM* (or *tarS*), the two fragments were then ligated by an overlap extension PCR by MA/MD (SA/SD) primers. pIMAY-Z_Δ*tarM* or pIMAY- Z_Δ*tarS* was obtained by setting a SLiCE reaction by 1 μL DNA insert, 1 μL linearized pIMAY-Z, 6 μL of H_2_O, 1 μL of SLiCE and 1 μL 10× ligation buffer at 37°C for 15 min. The plasmid was modified in *E. coli* IMO8B and was then electroporated into *S. aureus*. After incubation at 30°C for 48h on BHI agar containing 10µg/ml chloramphenicol and 100 µg/mL X-gal, transformants were selected and underwent steps of plasmid integration at 37°C and plasmid excision at 30°C. Finally, gene deletion was checked using primers MA and MD (or SA and SD).

TB4Δ*tarS* was constructed by transducing pIMAY-Z_Δ*tarS* plasmid to TB4 WT. Briefly, phage lysates of ɸ11Δ*int*-pIMAY-Z_Δ*tarS* was prepared by using ɸ11Δ*int* to infect TB4Δ*agrA*_pIMAY-Z_Δ*tarS* stored at 30°C following the above method of phage liquid infection. Afterwards, overnight cultures of TB4 strain were diluted and grown to OD_600_ of 1.4 before being added with 4.4 mM CaCl_2_. 1mL of TB4 cells together with 100 µL ɸ11Δ*int*- pIMAY-Z_Δ*tarS* lysates were incubated at 37°C for 20 min, following addition of 3mL TTA (6.0g TSB; 1.5g agar). Mixtures were then plated on TSA plates containing 17mM sodium citrate and 5 µg/mL erythromycin. Transductants were selected after 24h incubation at 37°C and the subsequent steps were carried out as described in SLiCE cloning method.

Complemented and overexpression mutants were constructed using expression vector pSK9067 (Brzoska & Firth, 2013). The primers MO and SO with *Sal*I and *EcoR*I restriction sites were used to clone *tarM* and *tarS* from TB4. The digested fragment was then ligated with the linearized pSK9067 harboring a P_spac_ promoter to construct the plasmid pSK9067_*tarM*/S, which was transformed into *E. coli* IM08B before electroporation into *S. aureus*. *S. aureus* transformants were selected on TSA containing 10µg/mL erythromycin.

The 8325-4 ɸ11Δ*int* mutant strain was constructed by allelic replacement of the up- and downstream flanking regions, using the primers described in Table S3. Cloning of the flanking regions into the pMAD vector was achieved using the SLiCE method as previously described, before transformation into strain 8325-4 ɸ11 (Zhang *et al*, 2014). Integration of the pMAD plasmid and crossover events were performed as previously described (Arnaud *et al*, 2004). Gene deletion was confirmed by the extraction of DNA from potential clones and PCR followed by sequencing using oligonucleotides that annealed outside the recombination flanks.

### IgG deposition assay

Anti-α-GlcNAc WTA IgG deposition was measured via binding of a secondary fluorescein 5- isothiocyanate (FITC) labeled Fab-Fragment against human IgG. Briefly, *S. aureus* strains were grown overnight at 37°C with agitation, then were subcultured with the treatment of 1 µM AIP-I (if necessary) and diluted to OD_600_0.4 in PBS+0.1% BSA, and the pellet was collected and resuspended in PBS+0.1% BSA. 25 µL of anti-α-GlcNAc WTA IgG in PBS+0.1% BSA prepared as described (Hendriks *et al*., 2021) were incubated with 25µL of *S. aureus* suspensions on ice before washing with PBS+0.1% BSA. Afterwards, the cells were incubated with F(ab’)2-Goat anti-Human IgG-FITC (Merck Goat anti-Human IgG, F(ab’)2, FITC Conjugated Affinity Purified, Catalog Number AQ112F; diluted 1:500) on ice, with the final anti-IgG concentration being 2.5µg/mL. The cells were washed once with PBS+0.1% BSA and fixed in 1% paraformaldehyde in PBS for 15min, followed by washing with PBS. The final suspensions in PBS were transferred to flow cytometry to measure fluorescence.

The normalization was done by setting the highest measured median fluorescence of all strains for each replicate to 100%. The WTA-specific Fab fragment 4461 as well as the B12 control Fab fragment were kindly provided by Prof. N. van Sorge (Amsterdam UMC, The Netherlands) (Hendriks *et al*., 2021).

### Co-culturing experiments

Bacteria were grown at 37°C, 180 rpm overnight and optical density (OD_600_) was measured before standardizing to the same. Overnight bacterial cultures were added to fresh TSB at 1:1000 dilution, followed by incubation at 37°C, 180 rpm overnight. Afterwards, 100 µL cultures were collected and subsequently used for susceptibility test against Stab20 by phage spot assay. Plate counting on TSA-X-gal (100 g/mL) was used to count colonies of each species in the co-cultures, as *S. pseudintermedius* and *S. haemolyticus* producing β-galactosidase generated blue colonies on the plate. *S. caprae* was differentiated on TSA-0.5% blood agar, which formed a hemolysis zone distinct from TB4.

### Statistical analysis

Statistical analysis was performed by using GraphPad Prism (GraphPad Software, version 9.4.0). Statistically significant differences were calculated by using one-way ANOVA method with Dunnett’s multiple comparison test. *P* values of <0.05 were considered significant (\*\*\**P* < 0.001; \*\**P*<0.01; \**P*< 0.05).

## Acknowledgements

We thank Prof. Mikael Skurnik of the Department of Bacteriology and Immunology at University of Helsinki for providing phage Stab20, and Prof. Friedrich Götz of the Interfaculty Institute for Microbiology and Infection Medicine (IMIT) at University of Tübingen for providing *S. pseudintermedius* strain ED99 and the mutant. Also we thank Dr. A. Robin Temming (Amsterdam UMC, the Netherlands) for the production of the WTA-specific and B12 control fragments; Prof. Christian A. Olsen (Department of Drug Design and Pharmacology, University of Copenhagen) for synthetic peptides and Assistant Prof. Nina M. Høyland-Kroghbo (Department of Plant and Environmental Science, University of Copenhagen) for valuable input to the work. J. Y. acknowledges the China Scholarship Council for financial support. A. P. acknowledges financial support from Deutsche Forschungsgemeinschaft, (SPP 2330 and PE 805/7-1); and infrastructural funding from the Cluster of Excellence EXC 2124 “Controlling Microbes to Fight Infections” project ID 390838134. H. I. and J. B. acknowledges financial support from the Independent Research Fund, Denmark, grant number 0135-00271B.

## Author contributions

**Jingxian Yang:** Conceptualization; data curation; formal analysis; investigation; methodology; visualization; writing–original draft; writing–review and editing. **Janine Zara Bowring:** Conceptualization; formal analysis; investigation; visualization; methodology; supervision; writing–review and editing. **Janes Krusche:** Formal analysis; investigation; **Benjamin Svejdal Bejder**: Methodology; **Stephanie Fulaz Silva:** Investigation; **Martin Saxtorph Bojer**: Methodology; **Tom Grunert:** Investigation; **Andreas Peschel:** Methodology; **Hanne Ingmer:** Conceptualization; formal analysis; visualization; supervision; funding acquisition; methodology; writing–review and editing.

